# A phylogeny of extant coleoid cephalopods with brain data

**DOI:** 10.1101/2024.04.29.591691

**Authors:** Kiran Basava, Theiss Bendixen, Alexander Leonhard Birk Sørensen, Nicole George, Zoé Vanhersecke, Joshua Omotosho, Jennifer Mather, Michael Muthukrishna

## Abstract

Extant coleoid cephalopods include over 800 species of octopuses, squid, and cuttlefish, which have drawn scientific and public interest for their complex behavior and cognition. Of these, approximately 10% (79) species have adult specimens with recorded measures of central nervous system size distributed across various sources. Here, we use a combination of topological placements from previous phylogenetic studies, along with mitochondrial and nuclear gene sequences obtained from GenBank, to build a composite phylogenetic tree with estimated branch lengths of all species with available brain measurements. This phylogeny is used for analyses in a forthcoming paper on cephalopod brain evolution, and ideally will be of use to other researchers interested in conducting comparative studies of coleoid cephalopod brains.

## Introduction

The origin of the subclass Coleoidea, containing today’s octopuses, squid, and cuttlefish, is dated to roughly 300 million years ago in the Devonian period, with the two major orders of Decapodiformes (containing squid and cuttlefish) and Octopodiformes (comprised of octopuses and the vampire squid) diverging in the mid-Triassic (Schweigert and Fuchs 2012; Tanner et al. 2017). Although these high-level splits have been established by fossil and genetic evidence, relationships within them including the placement of cuttlefish and bobtail squid, and the relationships between octopus species, have been more difficult to clarify (Lindgren & Anderson 2018; Sanchez et al. 2018).

In our paper ‘Coleoid Cephalopods Demonstrate Asocial Path to the Evolution of Big Brains’ (Basava & Bendixen et al. forthcoming), we test for correlates of brain evolution across coleoid cephalopods using a newly constructed trait database and the updated phylogeny described here. Often, phylogenetic analyses of trait evolution use a phylogeny broadly agreed upon by the field to conduct analyses, discarding taxa in the dataset which do not match those species on the phylogeny. For instance, a recent study analysing brain evolution in cephalopods by Ponte et al. (2021) includes 38 species in their phylogenetic analysis out of 78 in their total dataset, as these were the species that matched the cephalopod phylogeny from Lindgren et al. (2012). As we did not wish to exclude any taxa for which brain data were available from our analyses, we sought to build a phylogeny that would include all of these species.

### Methods

Rather than inferring a phylogeny from genetic data, we manually compiled a topology of species for which brain measurements were available based on the results of studies selected by Jan Strugnell, who lent her expertise as a cephalopod geneticist. We then added branch lengths to this topology using a phylogenetic inference of sequence data. This approach (manually compiling an informal topology from previously published results and adding branch lengths based on a constrained phylogenetic inference from genetic data) was taken on the advice of Dr. Strugnell based on the availability and accuracy of phylogenetic results and genetic data for coleoid cephalopods.

The starting point for this tree was the phylogeny from Lindgren et al. (2012), which is a relatively complete and widely-referenced phylogeny of cephalopods (López-Córdova et al. 2022; Sanchez et al. 2018, 2021; Strugnell et al. 2013). Relationships within clades were specified further using more recent phylogenetic analyses. Taxa were added in Newick format, either by hand or by using the ‘tree.merger’ function in the RRphylo package (Castiglione et al. 2018) depending on the number of new species added at a time. The Newick topology, as well as the placement for each species on the topology with the citation for the referenced study, are in https://github.com/kcbasava/ceph-brain-evolution/tree/main/phylogeny.

To add branch lengths, we conducted a topology-constrained phylogenetic inference from molecular data using BEAST 2.7.4 via the BEAUti GUI (Bouckaert et al. 2019). We constructed a dataset by downloading sequences for three nuclear genes: 18S rRNA (18S), 28S rRNA (28S), and rhodopsin (RHO), and three mitochondrial genes: 12S rRNA (12S), 16S rRNA (16S), and cytochrome oxidase I (COI) for all available species. Genetic data were not available for six species in our brain dataset; in those cases, sequences for closely related species (based on other studies using morphological characters) in the same genus were used^1^. The table containing GenBank accession numbers for all sequences used for each species is in https://github.com/kcbasava/ceph-brain-evolution/tree/main/phylogeny with notes indicating substituted species.

The genes were aligned separately using Multiple Sequence Comparison by Log-Expectation (MUSCLE) (Edgar 2004) with default settings as implemented on the NGPhylogeny.fr website (Lemoine et al. 2019). They were trimmed to obtain conserved blocks with Gblocks (Castresana 2000) likewise implemented on the NGPhylogeny.fr website using less restrictive settings allowing gaps in up to half the blocks (b1=50%, b2 = 85%, b3 = 10, b4 = 5, b5 =50%). Alignments were processed further to correct sequence names and add empty sequences for missing taxa for each gene in R using the packages seqinr (Charif et al. 2023) and ape (Paradis et al. 2019). Using BEAUti, the dataset was partitioned into noncoding and coding nuclear and mitochondrial genes, with four partitions total (COI; RHO; 18S and 28S; and 16S and 12S). Substitution models for the partitions were selected by bModelTest (Bouckaert & Drummond 2017). We set a Yule model of speciation as the tree prior and an optimized relaxed clock model (Douglas et al. 2021). The topology we compiled from previous phylogenetic studies described above was added as a multiple monophyletic constraint with substituted species names replaced. It contained multiple polytomies (non-binary speciation events where order of speciation is unclear) which were also resolved based on the BEAST analysis.

The divergence times were calibrated following the priors used by López-Córdova et al. (2022): for the root of the tree at the split between Decapodiformes and Octopodiformes using the Triassic coleoid fossil *Germanoteuthis* (Schweigert & Fuchs 2012) with a gamma distribution (offset=236, shape=2.0, scale=3.0, the split between Vampyromorpha and Octopoda using the Jurassic fossil *Loligosepia* (Fuchs & Weis 2008) with a gamma distribution (offset=195, shape=2.0, scale=3.0), the split between incirrate and cirrate octopods using the fossil *Styletoctopus annae* (Fuchs et al. 2009) with a gamma distribution (offset=93, shape=2.0, scale=3.0), and the origin of Loliginidae with the fossil *Loligo applegatei* (Clarke & Fitch 1979) with a gamma distribution (offset=48, shape=2.0, scale=3.0).

The MCMC was run for 30 million generations sampled every 3000 generations with a 10% burn-in. Tracer v1.7.2 (Rambaut et al. 2018) was used to ascertain effective sample sizes (>200) and chain convergence. The posterior distributions of trees from two independent BEAST runs were combined using LogCombiner (Bouckaert et al. 2019). A consensus tree (maximum clade credibility tree, highest product of posterior node probabilities) was also obtained using the software TreeAnnotator v2.7.4 (Fig. 2).

**Figure 1.**
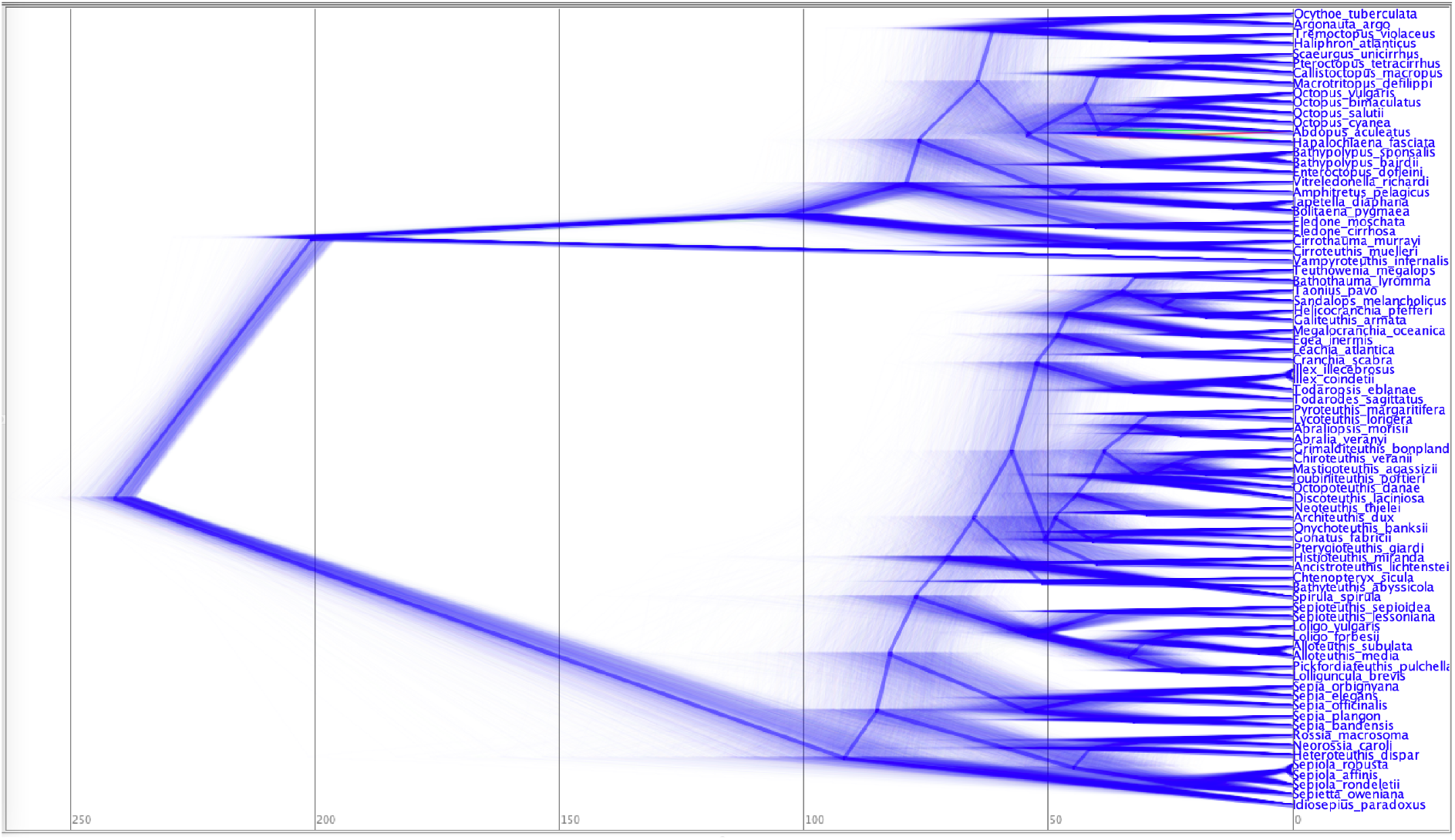
The posterior distribution of trees from the topology-constrained analysis in BEAST visualized with DensiTree v2.7.4 (Bouckaert & Helped 2014).

**Figure 2.**
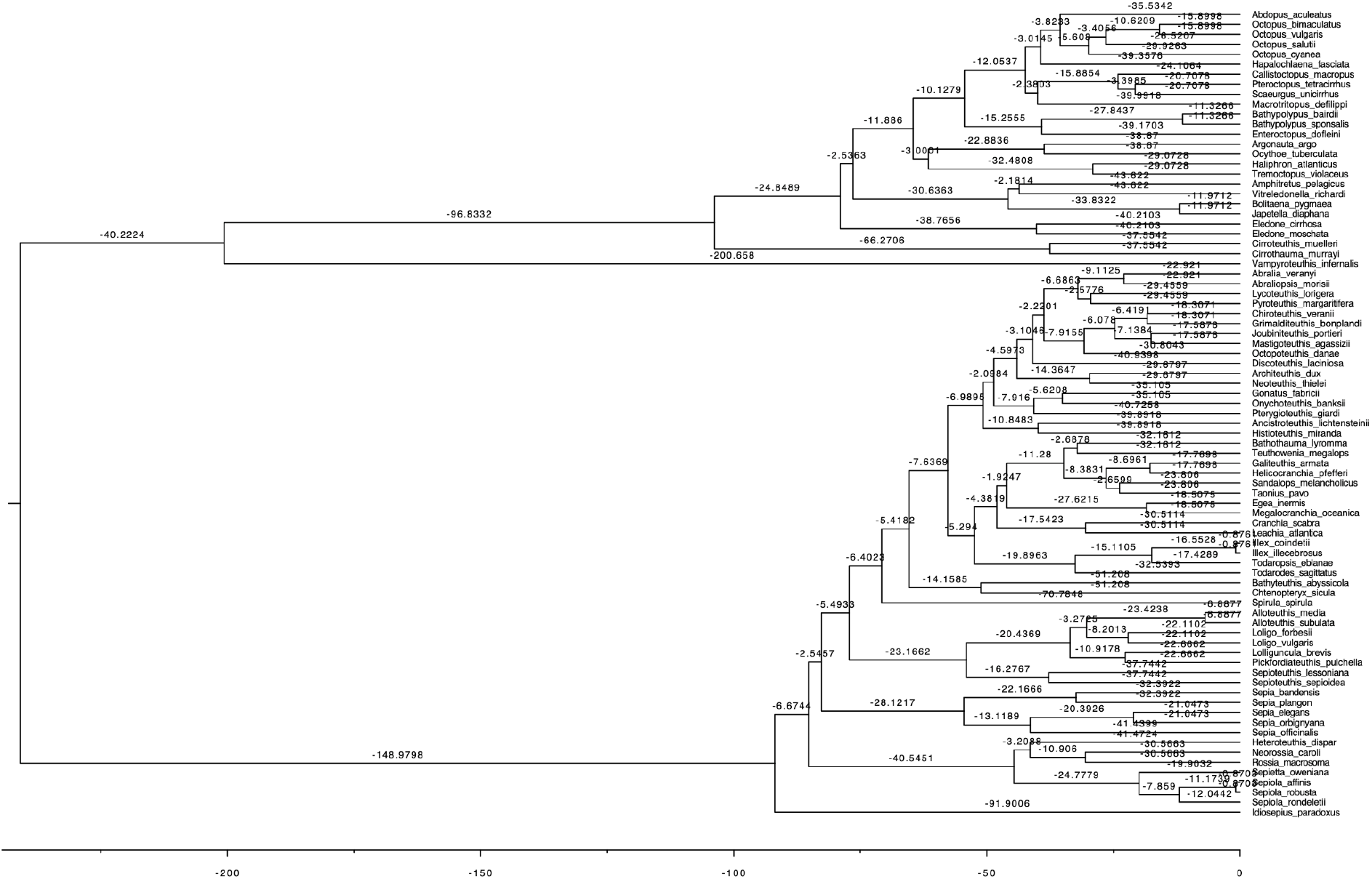
Estimated divergence times on consensus phylogeny using FigTree.

**Figure 3.**
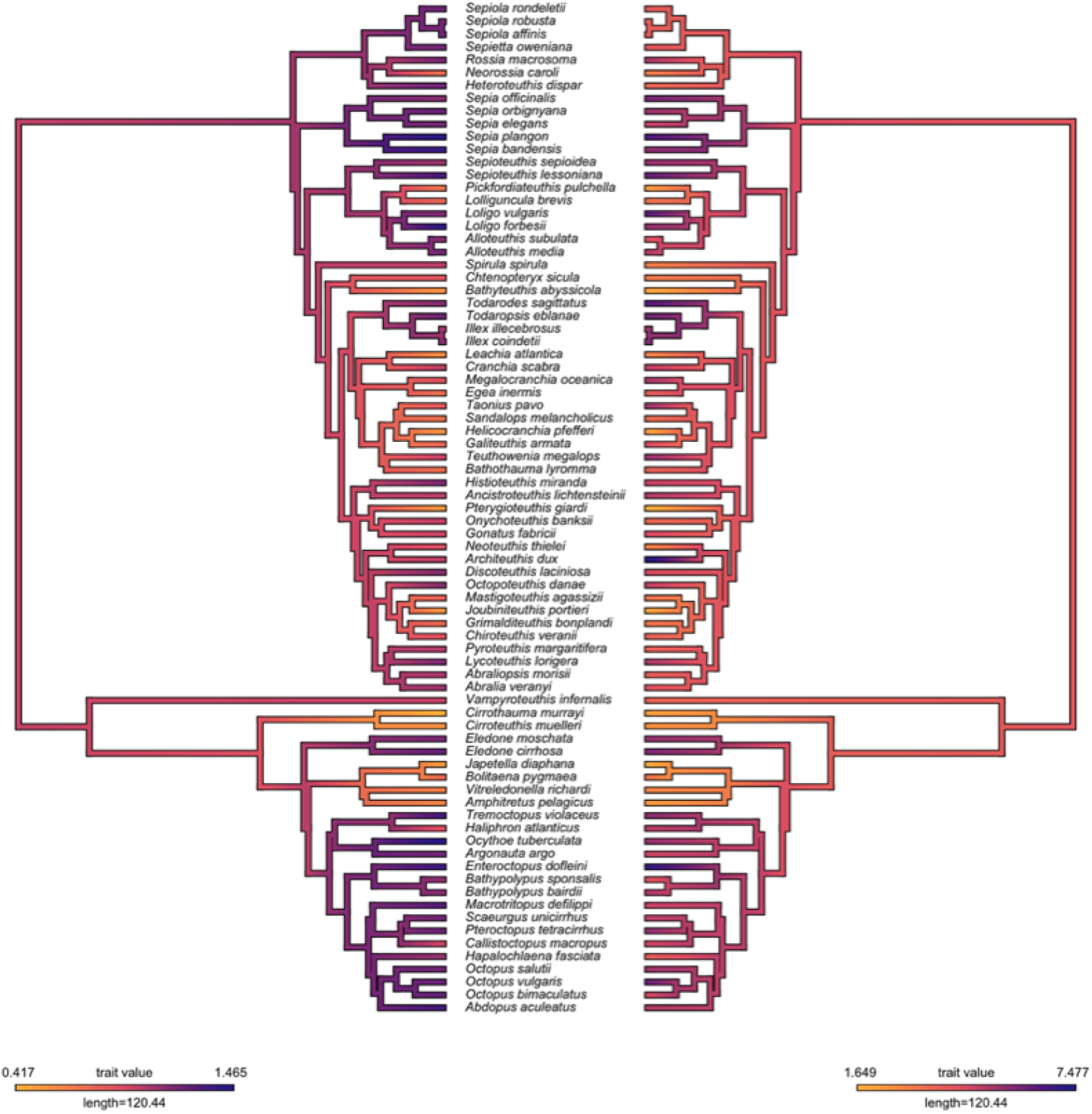
Ancestral state estimates for relative brain size (CNS/ML, left) and total brain size (CNS, right) on a log scale.

### Ancestral state reconstruction

The ancestral states of brain size (CNS) and relative brain size (CNS/ML) was estimated for the root and fossilized nodes using the anc.Bayes function from the phytools package in R (Revell 2024). This was estimated on the consensus phylogeny with the MCMC run for 100000 generations with a 20% burn-in.

## Results

Estimates for divergence times of major clades are shown in Table 1 with corresponding fossil calibrations. The origin for Decapodiformes (which was not fossil constrained) was estimated at 91.90 Mya (120.91 - 68.25) with *Idiosepius paradoxus* assigned as the first diverging group in the topological constraint.

**Table 1.**
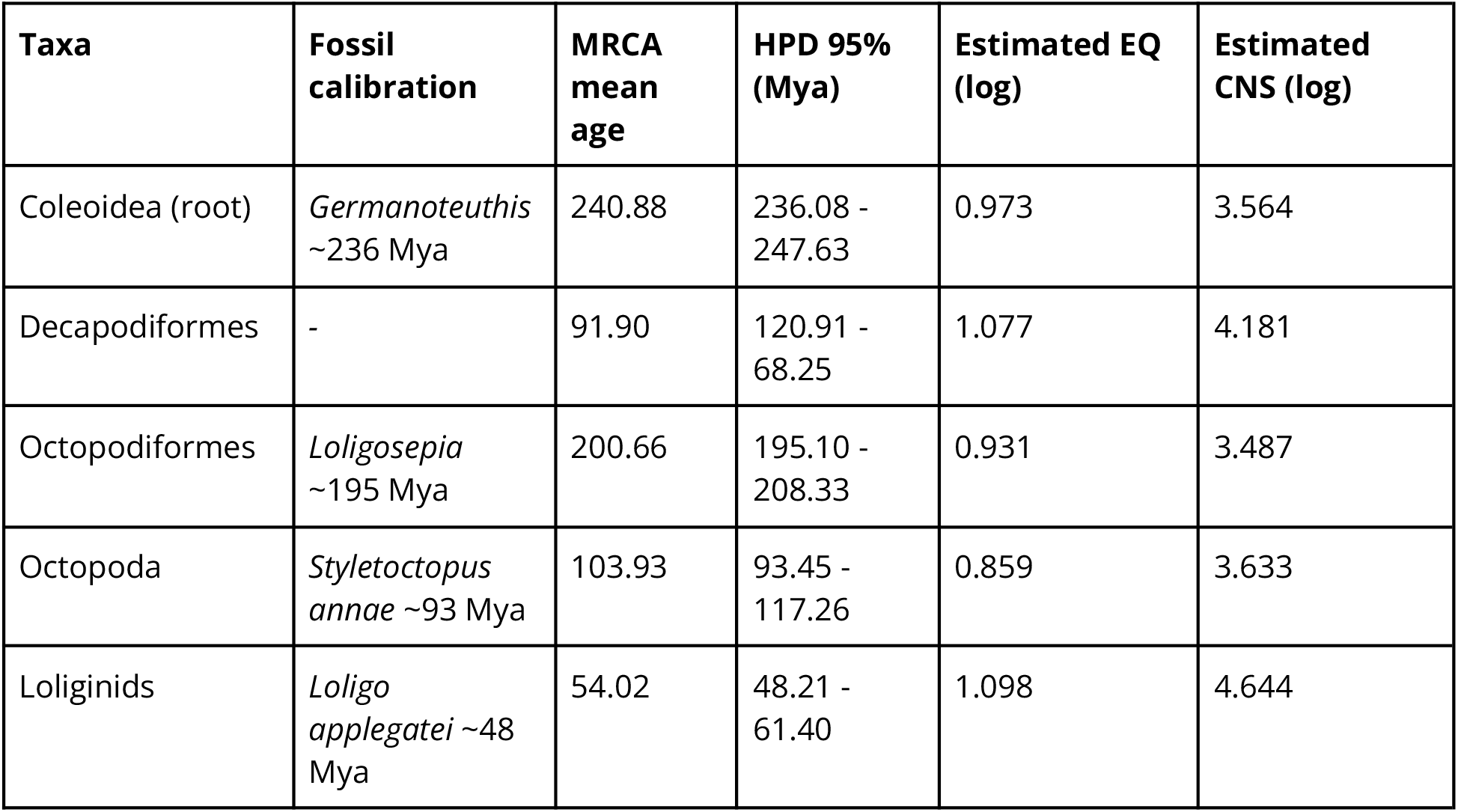
Estimated divergence times (95% highest posterior density interval) and means for fossil-calibrated nodes.

## Data

The XML files for the BEAST analyses, file containing GenBank accession numbers for each species, brain and body data, output posterior trees, and consensus tree are available at https://github.com/kcbasava/ceph-brain-evolution/.

## Acknowledgements

We thank the Templeton World Charity Foundation for supporting this work (Grant TWCF0464). We would also like to thank Jan Strugnell for assistance and advice in compiling and constructing the phylogeny.

## Footnotes

1 While some genera, e.g. Octopus, are likely paraphyletic and genus title is not a reliable indicator of sister relationships, this was not the case for any of the substituted species.

